# Molecular characterization of AMPA receptor trafficking vesicles

**DOI:** 10.1101/2021.06.09.447771

**Authors:** John Jacob Peters, Jeremy Leitz, Juan A Oses-Prieto, Alma L Burlingame, Axel T. Brunger

## Abstract

Regulated delivery of AMPA receptors (AMPARs) to the postsynaptic membrane is an essential step in synaptic strength modification, and in particular, long-term potentiation (LTP). While LTP has been extensively studied using electrophysiology and light microscopy, several questions regarding the molecular mechanisms of AMPAR delivery via trafficking vesicles remain outstanding, including the gross molecular make up of AMPAR trafficking organelles and identification and location of calcium sensors required for SNARE complex-dependent membrane fusion of such trafficking vesicles with the plasma membrane. Here, we isolated AMPAR trafficking vesicles (ATVs) from whole mouse brains via immunoprecipitation and characterized them using immunoelectron microscopy, immunoblotting, and liquid chromatography tandem mass spectrometry (LC-MS/MS). We identified several proteins on ATVs that were previously found to play a role in AMPAR trafficking, including SNARES (including synaptobrevin 2), Rabs, the SM protein Munc18-1, a calcium-sensor (synaptotagmin-1), as well as several new markers, including synaptophysin and synaptogyrin on ATV membranes. Additionally, we identified two populations of ATVs based on size and molecular composition: small-diameter, synaptobrevin-2- and GluA1-containing ATVs and larger transferrin-receptor-, GluA1-, GluA2-, GluA3-containing ATVs. The smaller population of ATVs likely represents a trafficking vesicle whose fusion is essential for LTP. These findings reveal the important role of AMPAR sorting into fusion-competent trafficking vesicles that are implicated in synaptic strength modification and reveal candidates of putative effectors and regulators of AMPAR trafficking.

## Introduction

At glutamatergic synapses, AMPARs are responsible for the largest component of postsynaptic responses in the form of cation influx, and along with NMDARs, are major contributors to various forms of synaptic plasticity including LTP (Dingledine et al., 1999; Malinow & Malenka 2002; Bredt & Nicholl 2003; Collinridge et al., 2004; Shepherd & Huganir 2007; Newpher & Ehlers 2008). Upon the arrival of an action potential, glutamate is released from synaptic vesicles into the synaptic cleft where it binds to postsynaptic AMPARs. When bound with glutamate, AMPARs open, allowing cations to enter and depolarize the postsynaptic cell. As a requisite of LTP (Malinow & Malenka 2002), the cellular correlate of memory (Nabavi et al., 2014), ATVs are exocytosed and AMPARs are recruited to the synapse, increasing the postsynaptic response (Lledo et al., 1998). The increased presence of AMPARs in the postsynaptic membrane has been characterized by light microscopy and electrophysiology studies, but little is known about the molecular composition of ATVs and the process by which they exocytose at the plasma membrane (Noel et al., 1999; Shi et al., 1999; Takumi et al., 1999; Liu et al., 2000; Passafaro et al., 2001; Ju et al., 2004).

AMPARs at the synapse come from two sources: receptors that have been recycled from the plasma membrane and receptors that have been synthesized *de novo*. Regardless of etiology, AMPARs are trafficked in AMPAR trafficking vesicles (ATVs) before they are inserted into the plasma membrane in a SNARE-dependent process (Jurado et al., 2013; Wu et al., 2017). While much is known about SNARE-dependent membrane fusion elsewhere in neurons (e.g., during neurotransmitter release via synaptic vesicle exocytosis), AMPAR insertion via ATV fusion has only recently begun to be elucidated. The insertion of AMPARs during LTP is particularly intriguing due to evidence that the process is calcium-triggered and involves synaptotagmins (Wu et al., 2017). Electrophysiology studies revealed that syntaxin 3 (Stx-3), SNAP-47, and synaptobrevin 2 (Syb2) are SNARE proteins involved in ATV fusion during LTP and that synaptotagmin-1 (Syt1) and −7 (Syt7) are the calcium sensors for this process (Jurado et al., 2013; Wu et al., 2017). Rab proteins, including Rab5, Rab8, Rab11, and Rab39, and the transferrin receptor (TfR) also play a key role in AMPAR delivery to synapses (Gerges et al., 2004; Liu et al., 2016). Despite these discoveries, there are many outstanding questions surrounding the ATV lifecycle, from ATV fusion to AMPAR endocytosis. For example, the cellular localization of most synaptotagmins is unknown. While Syt1, a key synaptotagmin involved in synaptic vesicle fusion, and other synaptotagmins have been found on synaptic vesicles, it is not known whether synaptotagmins are likewise trafficked on ATVs. Moreover, it is unclear to what extent proteins are sorted as AMPARs are endocytosed, stored in recycling endosomes, and inserted back into the postsynaptic membrane.

Due to their small size, relatively low abundance (compared to synaptic vesicles), and relative transience *in vivo*, ATVs have been challenging to study (Kittler & Moss 2006). Electron microscopy studies have yet to uncover convincing evidence of ATVs at the synapse perhaps because deliveries of AMPARs to the postsynaptic membrane often happen after induction of synaptic plasticity. The transience of AMPAR delivery and the difficulty of specifically targeting synapses that are undergoing plasticity with electron microscopy makes studying the molecular components involved in AMPAR trafficking *in situ* challenging. Advances in organelle isolation from synaptosomes have made it possible to faithfully isolate small organelles, specifically synaptic vesicles, for molecular characterization (Ahmed et al., 2013). To overcome the problems associated with studying AMPAR trafficking *in vivo*, we have adopted a similar strategy to specifically isolate ATVs from synaptosomes purified from whole mouse brains. Subcellular fractions were purified from neurons using multiple rounds of differential centrifugation, after which ATVs were enriched by immunoprecipitation using the GluA1 subunit of AMPARs, selected due to the role of GluA1-containing AMPARs in long-term potentiation (Shepherd & Huganir, 2007). ATVs were characterized using immunoblotting, LC-MS/MS, and immunoelectron microscopy. Here, we offer the first unbiased characterization of GluA1-containing ATVs. LC-MS/MS confirms several previously identified proteins found to be involved in AMPAR trafficking and identifies new markers. Immunoelectron microscopy reveals heterogenous populations of ATVs in terms of protein compositions and vesicle diameters. Combined, these data provide insights into the importance of AMPAR sorting for LTP and offer an unbiased candidate list of proteins potentially involved in diseases of the synapse.

## Results

### ATV isolation from whole mouse brains

To characterize the molecular composition of ATVs, synaptosomes were purified from whole brains of 6-12 P20 mice and hypoosmotically lysed to release their contents (Ahmed et al., 2013). The resulting lysis pellet (LP2), comprised of synaptosome contents, was flash frozen and stored at −80°C until used. GluA1-containing ATVs were purified from LP2 by immunoprecipitation with anti-GluA1 antibody (Fig. 1A). Antibody was allowed to bind overnight at 4°C and was subsequently bound to protein G paramagnetic beads before ATVs were gently eluted by competing with a peptide corresponding to the antibody epitope. Western blot analysis confirmed the presence of GluA1 in LP2 and the eluate (Figure 1B). Additionally, Western blot analysis confirmed the presence of VGLUT1, a marker of glutamatergic synaptic vesicles (a potential contaminate), in LP2 but not in the eluate. Negative stain electron micrographs revealed that the purification yielded vesicles with a diameter of 102.7 nm ± 50.8 nm (arithmetic mean; Fig. 1D), marking the first time ATVs have been definitively visualized (Fig. 1C). To further confirm the fidelity of ATV purification, the same immunoisolation protocol was performed using LP2 purified from *GLUA1-/-* knockout mice. Western blot analysis confirmed the deletion of *GLUA1* but the retention of VGLUT1 expression. Additionally, no small vesicles were identified in the anti-GluA1 immuno-isolated sample with electron microscopy (Fig. 1E).

**Figure 1.**
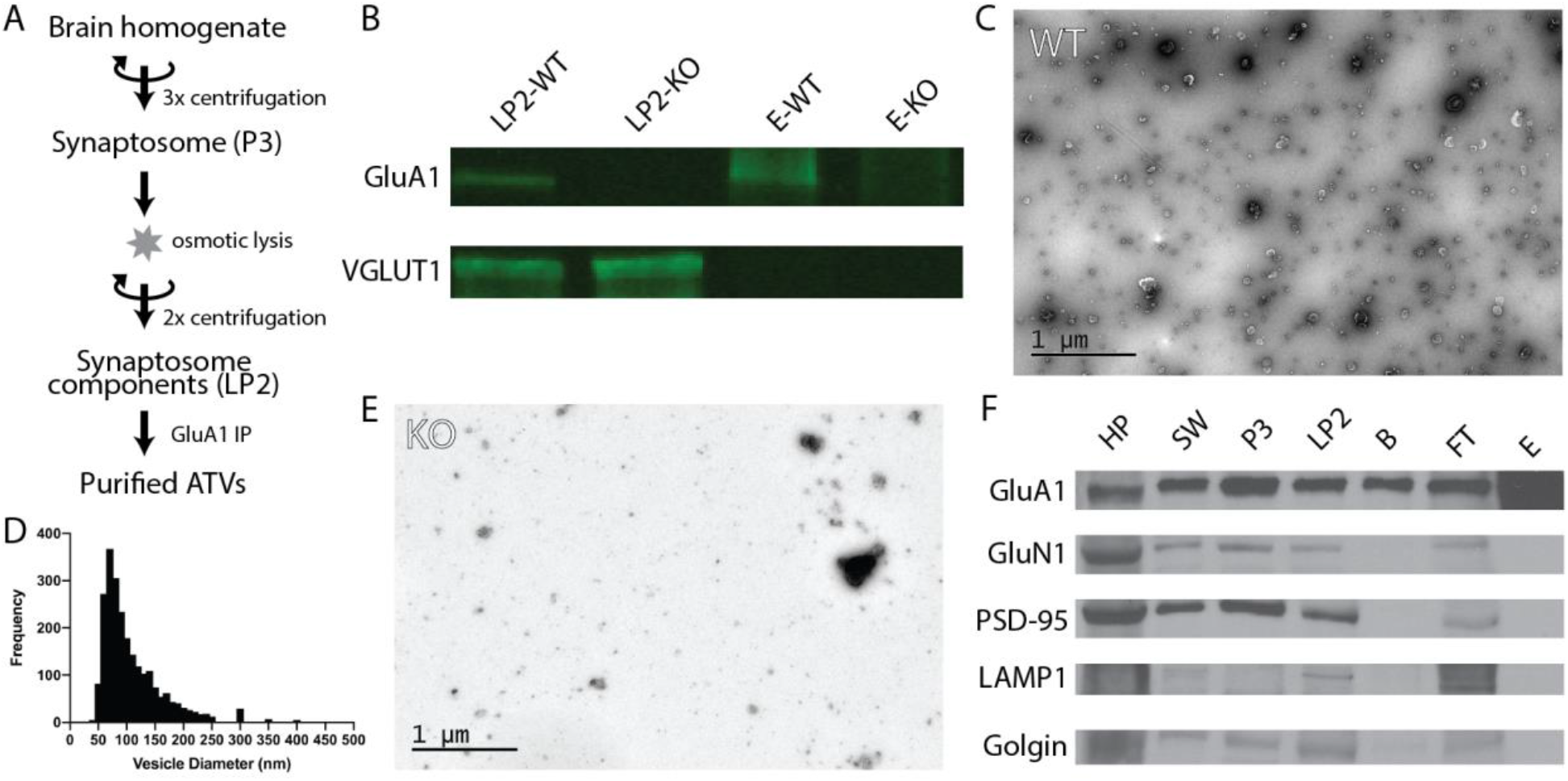
Purification of ATVs from whole mouse brain **(A)** Purification protocol for isolating ATVs. **(B)** Western blots for GluA1, VGLUT1, and GluN1 in WT isolated synaptosome content (LP2-WT), GluA1 KO isolated synaptosome content (LP2-KO), WT ATV eluate (E-WT), and GluA1 KO ATV eluate (E-KO). **(C)** Negative stain electron micrograph showing isolated ATVs. **(D)** Histogram of vesicle diameters. **(E)** Negative stain electron micrograph showing limited material in purification from GluA1 KO mice. **(F)** Western blots for GluA1, GluN1, PSD-95, LAMP1, and Golgin for homogenized whole brain pellet (HP), the second synaptosome wash step (SW), synaptosome (P3), synaptosome content (LP2), beads from immunoprecipitation prior to elution (B), flowthrough from immunoprecipitation (FT), and eluate off beads (E)

### Immunoisolation leads to pure ATVs

While initial results were suggestive of a relatively pure population of ATVs, we probed several additional molecules to further confirm eluate quality. Western blots were performed on samples from each step of the isolation process to monitor which molecular components were enriched (Fig. 1F). Confirming previous results, the GluA1 subunit of the AMPAR was identified throughout the isolation process and was enriched in the final eluate. Several other proteins were probed to verify isolation purity, including GluN1, PSD-95, LAMP1, and golgin. GluN1 is an NMDA receptor subunit and is also present in the glutamatergic postsynaptic compartment (Paoletti et al., 2013). Similarly, PSD-95 is a component of the postsynaptic density at excitatory synapses (Craven & Bredt, 1998). LAMP1 is a lysosomal marker (Griffiths et al., 1988), and golgin is a Golgi aparatus marker (Munro, 2011). All of these markers were identified in each step up until the immunoprecipitation and final elution, indicating that as expected, subcellular compartments were maintained throughout the preparation but were excluded during the immunoisolation.

### Immunoelectron microscopy revealed molecular components of ATVs

Immunoelectron microscopy was performed on ATVs to assess the frequency of protein localization on ATVs for several known AMPAR-associated proteins (Fig. 2A-G). Secondary antibody concentration was optimized to minimize non-specific, background gold (<1 free gold per field of view). A positive hit was defined as a gold particle within 5 nm of an ATV. AMPAR subunits GluA2 and GluA3 were probed to test for the presence of these subunits in the GluA1-affinity purified ATVs. GluA2 was found on 42.6% of ATVs, and the GluA3 subunit was found on 36.7% of ATVs. TfR, a known marker of AMPAR endosomes, was identified on 48.2% of ATVs. Synaptophysin1 (Syp1) was identified on 90.2% of vesicles. Syb2 was found 82.1% of ATVs, while Syt1 was identified on 44.0% of ATVs. Additionally, the diameters of ATVs that were labelled by GluA2 (140.1 ± 52.5 nm), GluA3 (134.5 ± 60.2 nm), TfR (121.7 ± 66.4 nm), Syp1 (116.2 ± 50.8 nm), Syb2 (93.9 ± 43.8 nm), and Syt1 (105.2 ± 54.6 nm) were measured (all arithmetic means; Fig. 2H). As a negative control, VGLUT1 (vesicular glutamate transporter), a marker of glutamatergic synaptic vesicles, was probed (data not shown), and only 8.4% of ATVs were positive for VGLUT1. The Kolmogorov-Smirnov test was performed, comparing the cumulative frequency distribution for each marker to the overall population of ATVs obtained from the negative stain experiments shown in Figure 1D (Fig. 2I). The cumulative frequency distribution for Syb2-labelled ATVs was significantly shifted to the left, indicating smaller diameters (p=0.0101), while the Syp (p<0.0001), TfR (p=0.0100), GluA2 (p<0.0001), and GluA3 (p<0.0001) distributions were significantly shifted to the right (larger diameters). Syt1 was not significantly shifted from the global ATV diameter distribution (p=0.9382). Smaller, Syb2-labelled vesicles are unlikely to be synaptic vesicles due to the low frequency of VGLUT1-labelled vesicles and the substantial difference in size between Syb2-labelled vesicles and the 40-45 nm diameter that has previously been reported for synaptic vesicles (Takamori et al., 2006). Additionally, the mean diameter of VGLUT-1 labelled vesicles (arithmetic mean of 97.6 ± 54.8 nm) is also much greater than the reported diameter of synaptic vesicles, which suggests that VGLUT1-labelled vesicles are most likely small endosomes or membrane fragments (Supplemental Figure 1).

**Figure 2.**
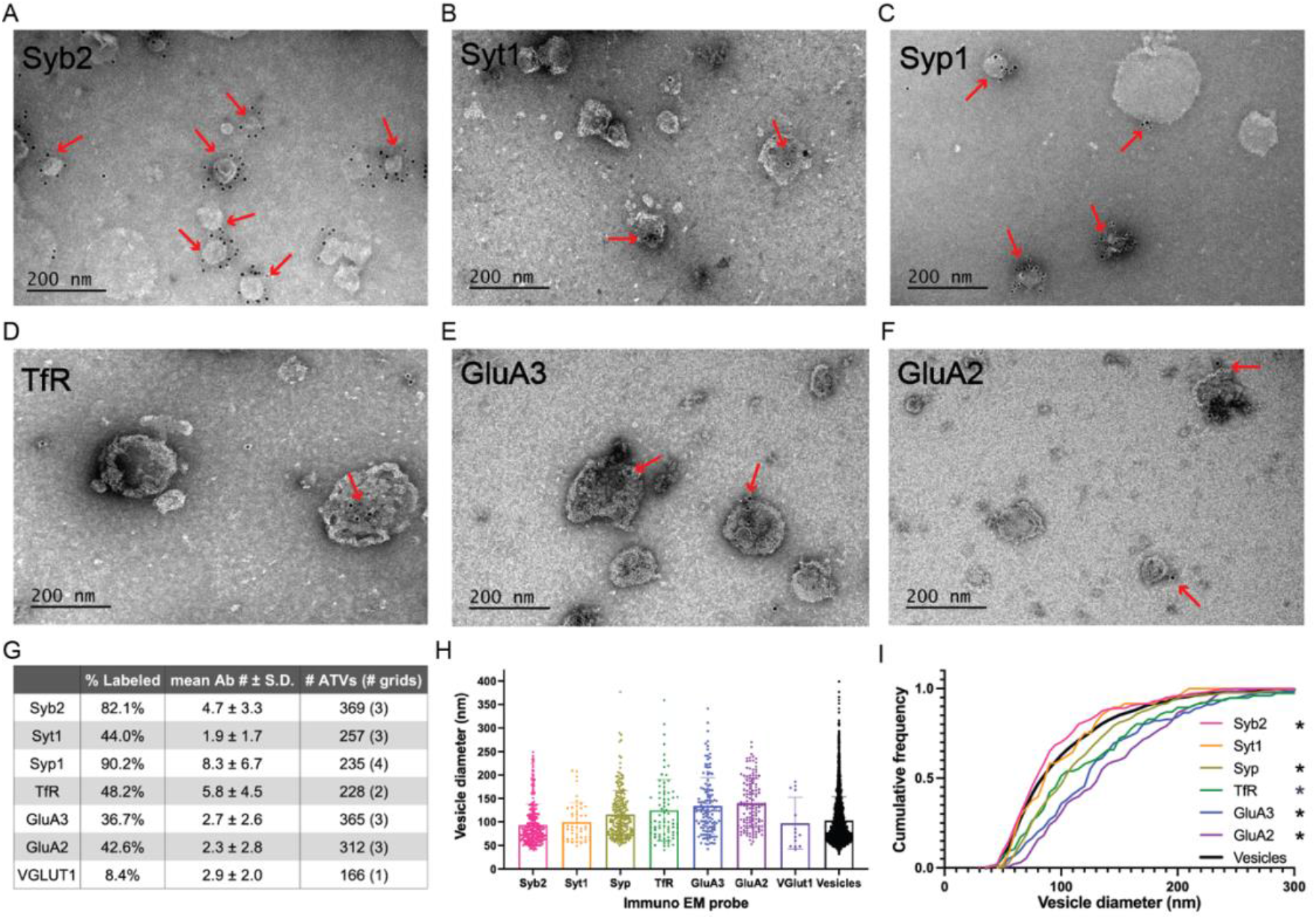
Western blot and electron microscopy analysis of purified vesicles. **(A-F)** Immunoelectron micrographs for GluA2, GluA3, Syb2, Syt1, TfR, and Syp1. **(G)** Summary table of immunoelectron microscopy analysis. **(H)** Mean and standard deviations of diameters of vesicles labelled with each probe during immuno EM. Individual histograms for each label are in Supplemental Figure 1. **(I)** Normalized cumulative frequency distributions of diameters of ATVs labelled with each probe during immuno EM. The bold line represents the frequency distribution of all vesicles from Figure 1D. The Kolmogorov-Smirnov test was performed to test statistical significance between an independent population of vesicles from Figure 1D and vesicles containing Syb2 (p=0.0101), Syt1 (p=0.9382), Syp (p<0.0001), TfR (p=0.0100), GluA2 (p<0.0001), and GluA3 (p<0.0001). * indicates p-value < 0.05.

### LC-MS/MS analysis identifies known and several new AMPAR trafficking proteins

LC-MS/MS was performed on isolated ATVs. We identified a total of 755 unique proteins with expectation values < 0.005 across three biological replicates (Fenyö & Beavis, 2003). We applied two additional filters to these 755 proteins to ensure high quality and abundance. Of those 755 unique proteins, 442 proteins were identified in two or more data sets (Fig. 3A). The sequence coverage (fraction of protein sequence that was identified) for 180 proteins was greater than 7.5%, suggestive of higher abundance. Proteins were manually categorized based on function and cellular localization (Fig. 3B). Cytosolic proteins, channels/transporters, and Rabs were the most commonly identified protein classes with 39, 23, and 21 hits respectively. Among the top proteins enriched in ATVs (Table 1) are AMPAR subunits GluA1, GluA2, and GluA3, as well as AMPAR-associated Dnajc13 (Perrett et al., 2015), TfR (Liu et al., 2016), neuroplastin (Jiang et al., 2021), and ABHD6 (Wei et al., 2016). In addition, the genes for Rab5, 8, 11, and 39, all implicated in AMPAR trafficking, were also among the top 180 candidates (Gerges et al., 2004). Furthermore, other synaptic proteins that have yet to be identified as AMPAR-trafficking-associated, including Syp1, synaptogyrin-1 (Syngr1) and −3 (Syngr3), and Munc18-1, were identified (Table 1).

**Figure 3.**
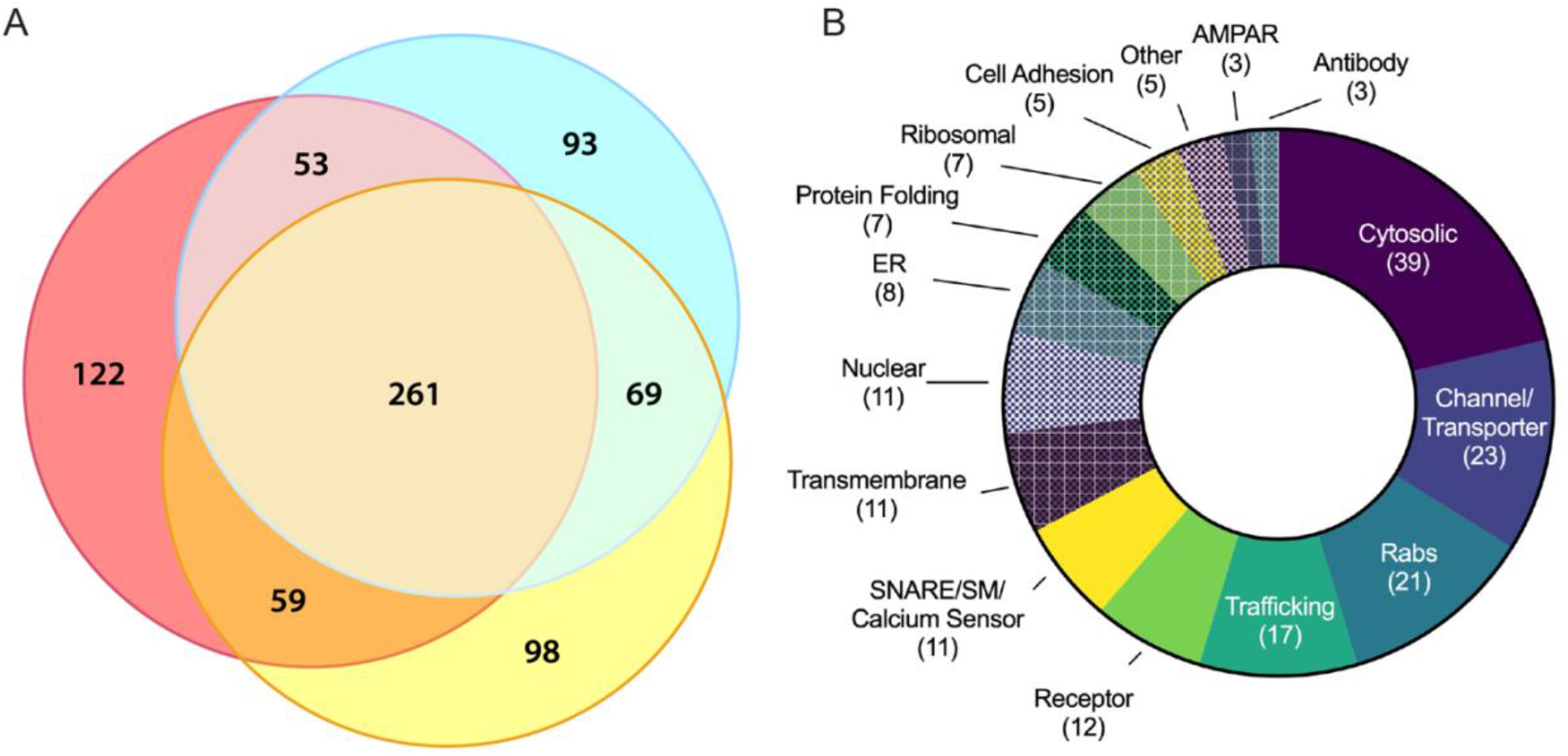
Molecular characterization of ATV proteins using LC-MS/MS **(A)** Three-way Venn diagram showing protein hits in three LC-MS/MS biological replicates with each color representing a biological replicate. **(B)** Protein ontology of identified candidates using gene ontology resource.

**Table 1.**
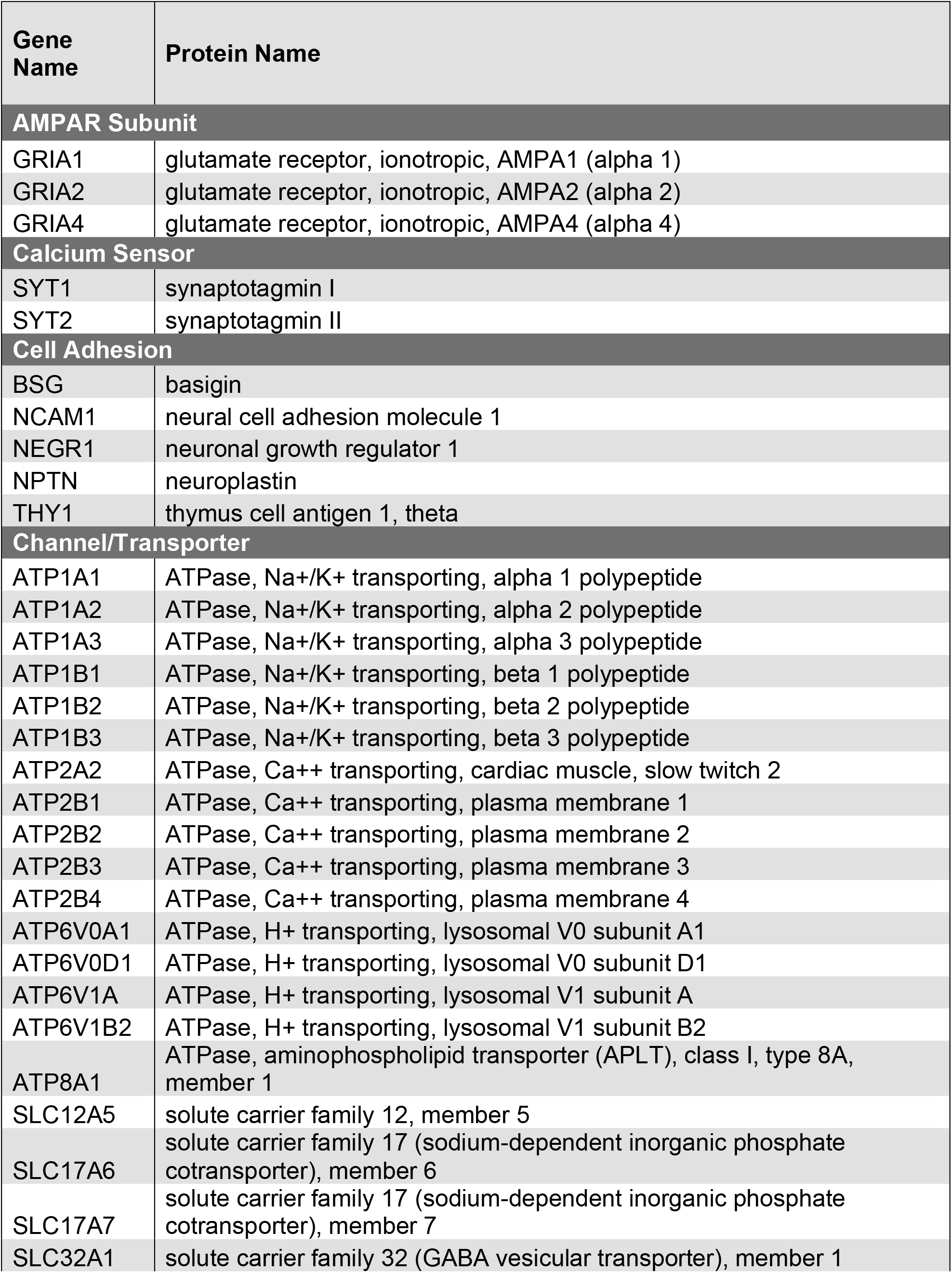
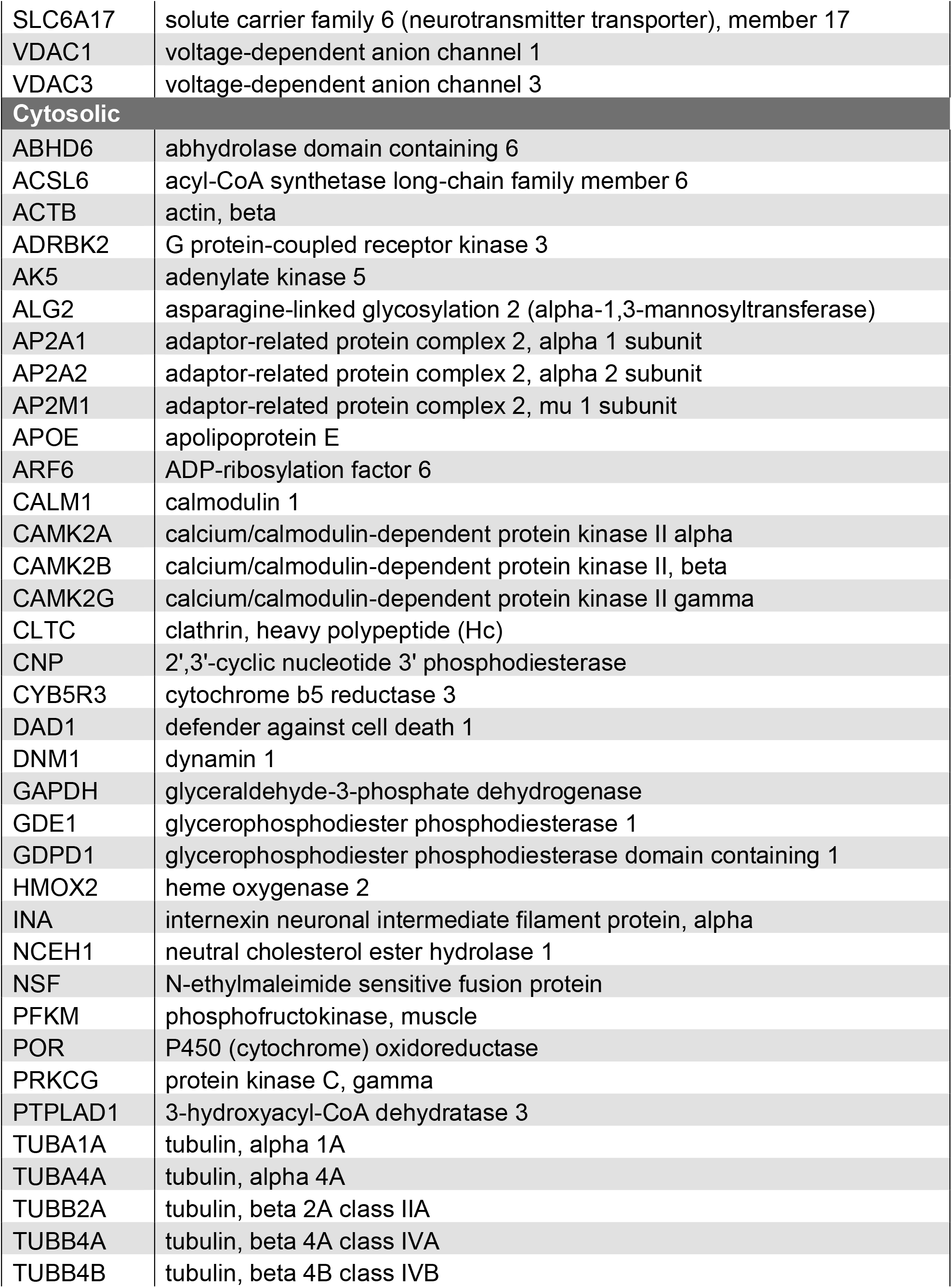
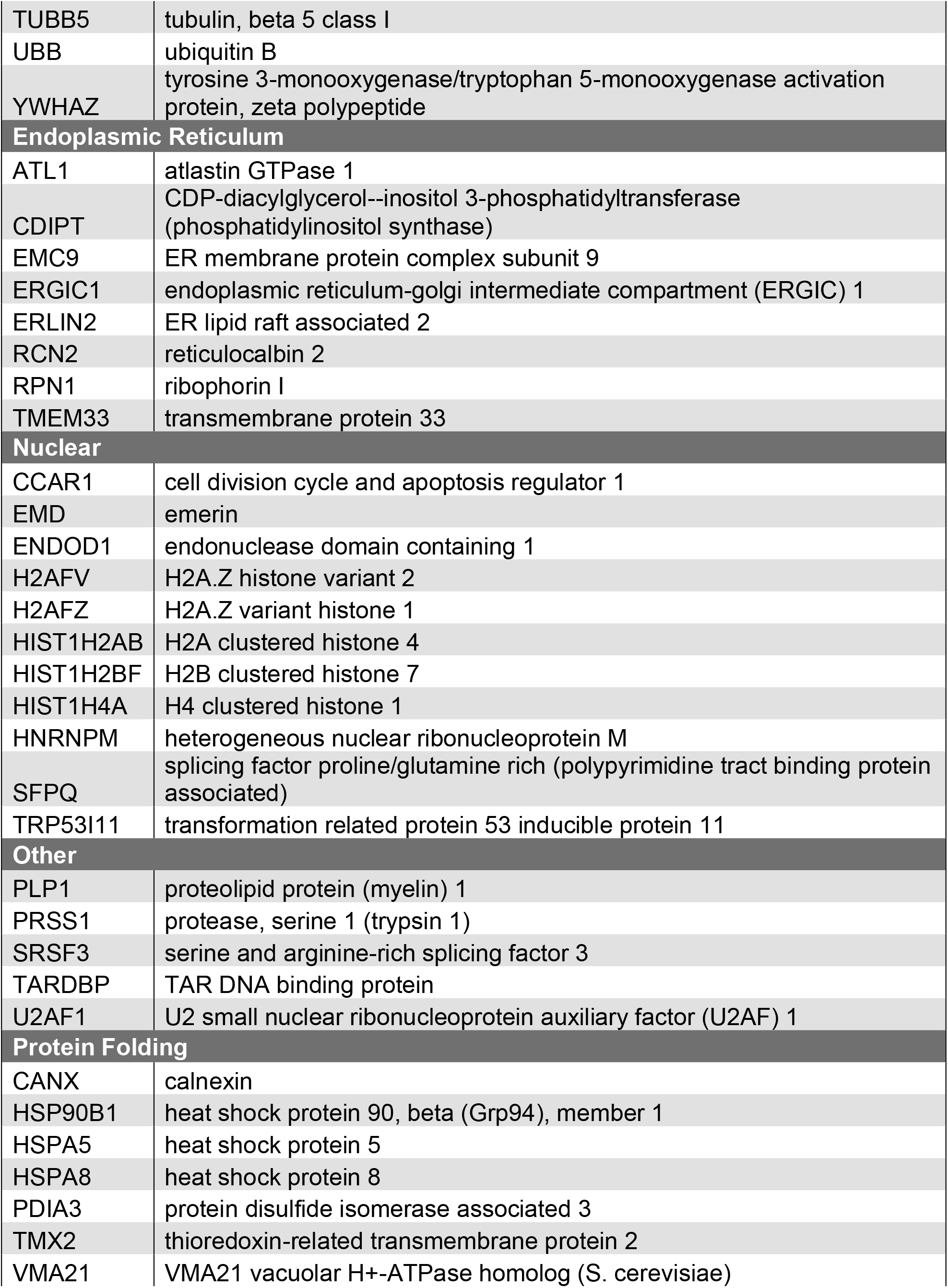
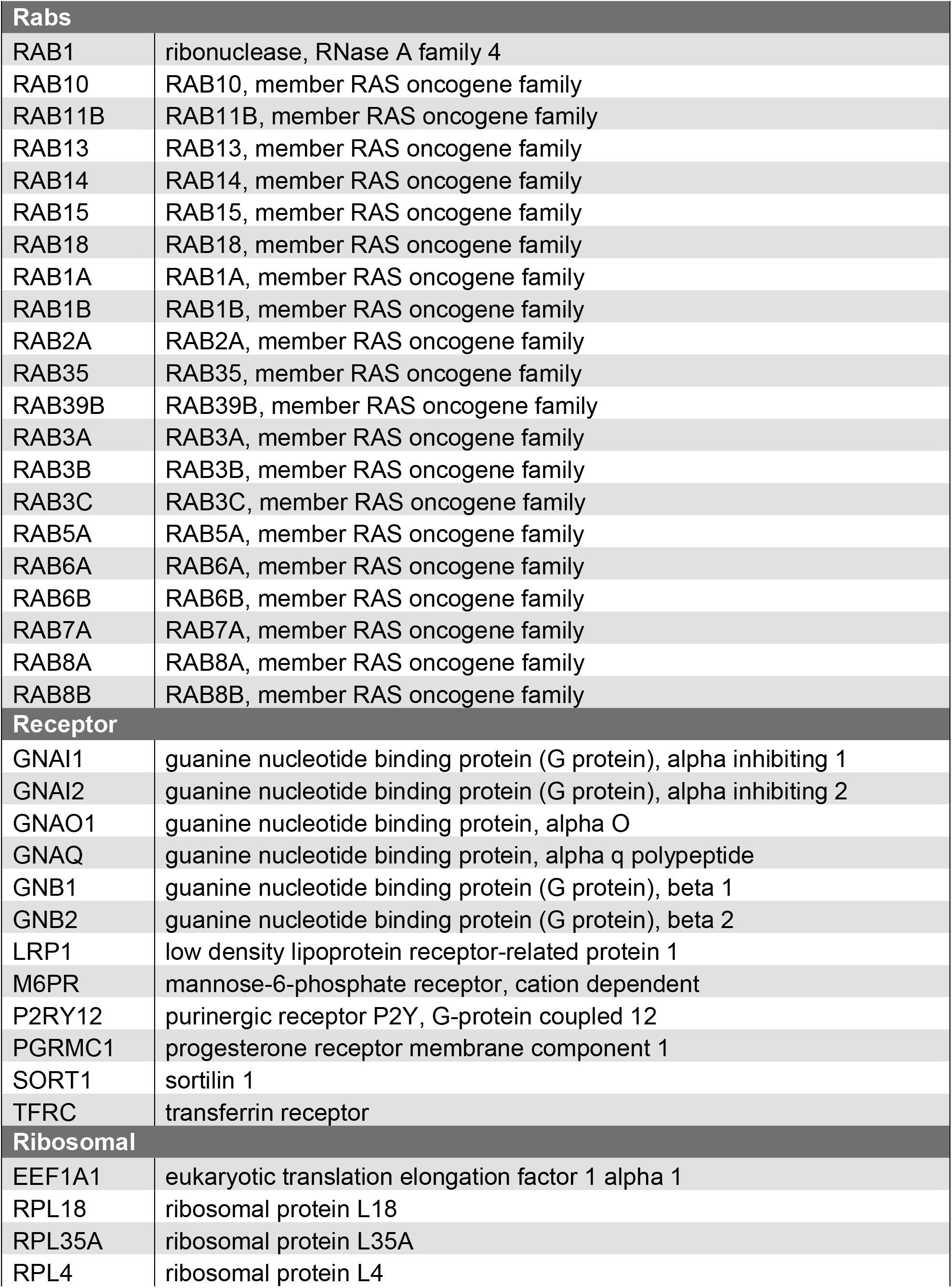
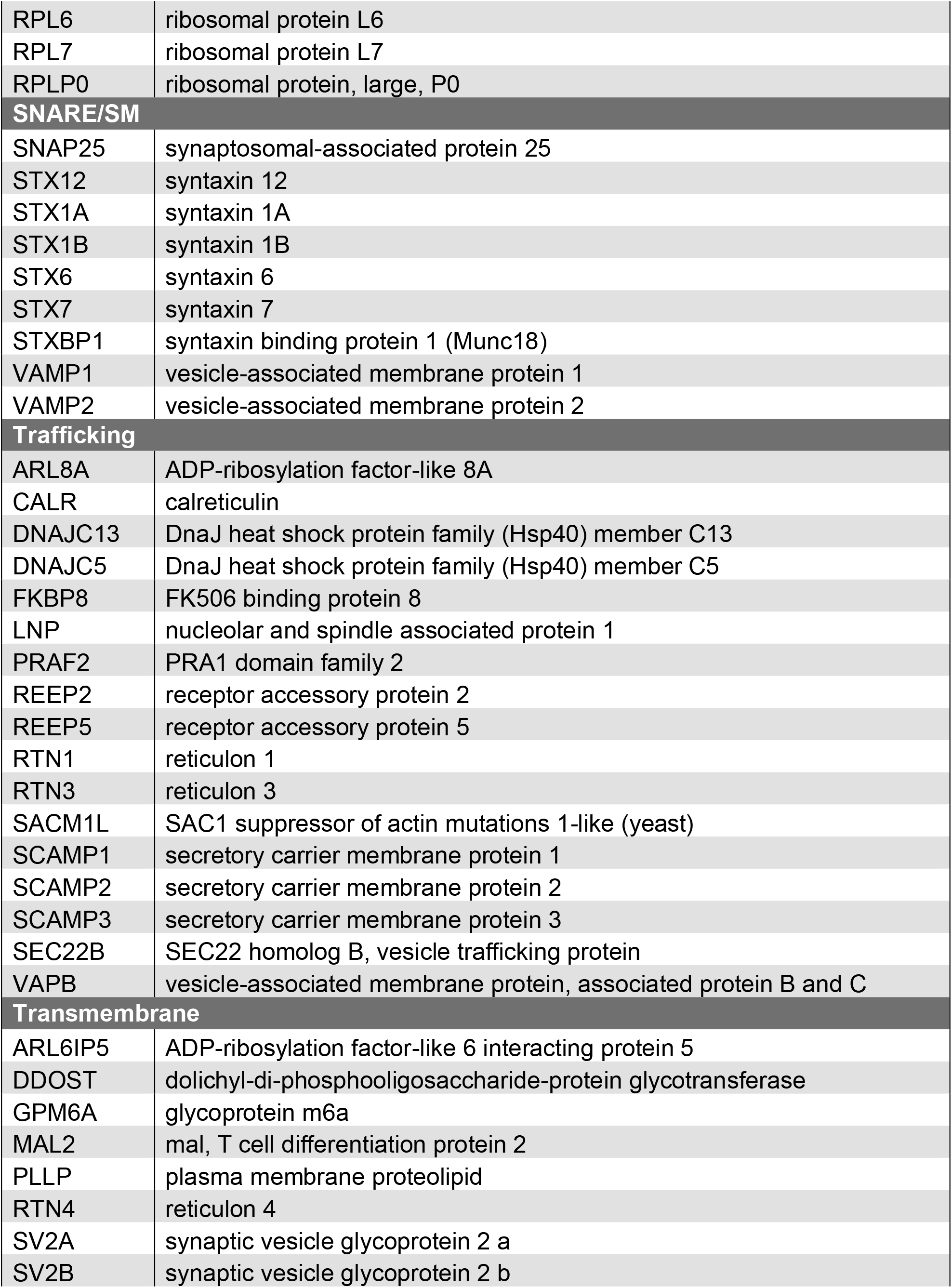
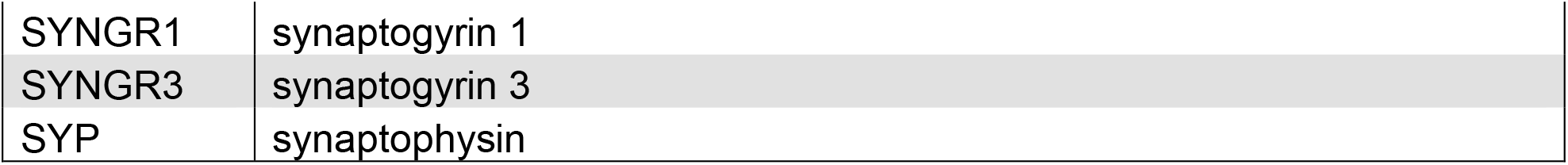
Protein ontology of top candidates identified with LC-MS/MS using gene ontology resource.

## Discussion

### ATVs can be specifically purified from whole mouse brains

Due to their relatively low abundance at synapses compared to other synaptic content (e.g., synaptic vesicles), ATVs have been difficult to characterize in the past. Here, we developed a protocol to specifically purify and enrich ATVs from synaptosome lysate via immunoprecipitation using a monoclonal anti-GluA1 antibody. Attempts to immunoprecipitate from GluA1 KO mice yielded little material, indicating that the purification process is specific. Western blot analysis of samples taken from steps in the purification process further supports the specificity of this isolation. Seven cellular components were probed by Western blot (Figure 2A). GluA1, the AMPAR subunit being enriched, was present in each step of the purification process and was enriched in the final elution. In contrast, GluN1 (NMDA receptor subunit), PSD-95 (postsynaptic density component), LAMP1 (late endosome component), and golgin (Golgi marker) were all present throughout the purification process but did not bind to beads nor appear in the final eluate. Typically, synaptosomes generated via differential centrifugation have primarily been used to study presynaptic components; our data indicate that synaptosomes present in the crude synaptosome fraction (P3) also preserve some postsynaptic material.

Immunoelectron microscopy analysis further confirmed the specificity of ATV purification. Unsurprisingly, Syb2, a SNARE protein essential for AMPAR insertion during LTP, labelled 82.1% of ATVs (Jurado et al., 2013). In addition, 42.6% of ATVs were positive for the GluA2 subunit of the AMPAR. This aligns well with evidence that GluA1/GluA2 heteromers are the most common AMPAR composition (Lu et al., 2009; Zhao et al., 2019). Furthermore, 36.7% of ATVs were positive for the GluA3 subunit. This could perhaps be reflective of GluA1/A3 heteromers; it has been previously observed that ~10% of GluA3-containing AMPARs also contain GluA1 (Wenthold et al., 1996; Diering & Huganir, 2018). Alternatively, multiple AMPARs could be contained in the same ATV, and this observation could be reflective of GluA2/A3 heteromers.

LC-MS/MS also provided supportive evidence that ATV purification is specific. The GluA1, GluA2, and GluA4 subunits were all identified in the top mass spectrometry hits. GluA3 was also identified but had lower sequence coverage, possibly due to sequence similarity between it and other AMPAR subunits. Several of the top hits identified via mass spectrometry were Rab proteins, including Rab5, Rab8, Rab11, and Rab39, which have been previously characterized as proteins required for AMPAR trafficking (Gerges et al., 2004). Rab39 contributes to AMPAR trafficking from the endoplasmic reticulum to the Golgi, and mutations in this protein have been connected to autism spectrum disorders (Mignogna et al., 2015). Rab5 is required for AMPAR endocytosis (Brown et al., 2005), while Rab8 and Rab11 (Brown et al., 2007) have been implicated in AMPAR insertion into the plasma membrane. Mass spectrometry also identified several other proteins associated with AMPAR trafficking, including Lrp1 (Gan et al., 2014), TfR (Liu et al., 2016), Dnajc13 (Perrett et al., 2015), TDP-43 (Schwenk et al., 2016), and Abhd6 (Wei et al., 2016). The immunoelectron microscopy and mass spectrometry data combined support the conclusion that ATVs can reliably be purified from synaptosomes, generated from homogenized, whole brain tissue. The enrichment and concentration of ATVs allowed for new access to further characterize the molecular components associated with AMPAR delivery to the synapse and insertion in the postsynaptic membrane, and the molecular characterization of ATVs presented here is the first time ATVs have been isolated and characterized in a high-throughput manner.

### Insights into ATV lifecycle and AMPAR insertion

One outstanding question in the AMPAR lifecycle is to what degree proteins on ATVs are sorted as AMPARs are endocytosed, travel to recycling endosomes, and are reinserted back into the postsynaptic membrane. Immunoelectron microscopy combined with vesicle diameter analysis identified at least two possible unique populations of ATVs. Specifically, the cumulative frequency distribution of the average diameters of ATVs labelled with TfR was significantly shifted to the right compared to an independent overall population of ATVs, while the cumulative frequency distribution of the average diameters of ATVs labelled with Syb2 was significantly shifted to the left. These right-shifted, or larger, ATVs were also more likely to contain GluA2 and GluA3. Thus, purification of ATVs from whole mouse brain isolates at least two populations of ATVs (Figure 2H-I). Larger ATVs containing TfR and a mixed population of AMPAR subunits may represent recycling endosomes, while smaller ATVs with fusion machinery may represent either *de novo* ATVs containing newly synthesized AMPARs or ATVs that have been formed from recycling endosomes. The identification of multiple populations of ATVs reveals key protein sorting that occurs as a requisite for AMPAR delivery during LTP.

Many of the SNARE and SNARE effector proteins involved in AMPAR insertion during LTP have been identified, including Stx-3, SNAP-47, and Syb2 (Jurado et al., 2013). Additionally, the N-terminal, Sec1/Munc18-like-binding portion of Stx-3 is essential for LTP (Jurado et al., 2013), providing evidence for the possible role of Munc18 in AMPAR insertion. Munc18-1 is associated with ATVs as observed by mass spectrometry, and combined with evidence that Munc18-1 binds to Stx-3 (Hata & Südhof, 1995), Munc18-1 is a likely candidate for a regulator of AMPAR insertion. In synaptic vesicle fusion, Munc18 stabilizes syntaxin-1A (Südhof, 2013), and Munc13 is required to aid in the transition of the syntaxin/Munc18 complex into the ternary trans-SNARE complex, a critical step to ensure parallel assembly of all SNARE complex components (Ma et al., 2013; Lai et al., 2017; Brunger et al., 2019). After fusion, the ternary SNARE complex is disassembled with the ATPase, NSF, and adaptor protein, SNAP, for use in future fusion events (Sollner et al., 1993; Mayer et al., 1996; Hanson et al., 1997). Therefore, Munc18, Munc13, NSF, and SNAP could also play roles in regulating SNARE assembly and disassembly during ATV fusion. Additionally, LC-MS/MS identified synaptotagmin-2 (Syt2), a calcium sensor that performs equivalent functions to Syt1 (Pang et al., 2006), as a marker of ATVs. Indeed, only 44.0% of ATVs contained Syt1 (Fig. 2G), consistent with the implication of alternative calcium sensors such as Syt2 or Syt7 for AMPAR insertion (Wu et al., 2017). Furthermore, it is worth noting that the exact location of AMPAR insertion is an active area of exploration (Choquet and Hosy, 2020). Our results are agnostic to precise fusion location, compatible with ATV fusion happening either perisynaptically or directly at the synapse. Future studies should explore the localization and roles of these SNARE effector proteins in AMPAR insertion.

### New ATV trafficking candidates

Mass spectrometry and immunoelectron microscopy revealed several new candidates with connections to AMPAR trafficking and neurological disease. Syp1, best known as a synaptic vesicle marker, densely labeled ATVs. Interestingly, Syp1, Syngr1, and Syngr3 were identified among the top mass spectrometry hits. Previously, synaptophysin and synaptogyrin have been shown to cooperatively contribute to LTP (Janz et al., 1999). Furthermore, Syngr3 has been implicated in tauopathies, and the reduction of Syngr3 expression in neurons has been shown to rescue synaptic plasticity deficits induced by tau (Largo-Barrientos et al., 2021). While important roles for synaptophysin and synaptogyrin have already been identified in the presynaptic terminal, the potential for a postsynaptic contribution has yet to be explored.

### Connections to disease

Many of the candidates identified in mass spectrometry have been implicated in neurological disorders. The knockdown of TDP-43, a protein implicated in amyotrophic laterals sclerosis (ALS) (Sreedharan et al., 2008), decreases the number and motility of Rab-11 endosomes which in turn impairs AMPAR recycling (Schwenk et al., 2016; Esteves da Silva et al., 2015). LRP1, previously implicated in both Alzheimer’s disease and GluA1 trafficking, was also identified by mass spectrometry (Liu et al., 2010; Gan et al., 2014). Finally, Dnajc13, a known contributor to Parkinson’s disease (Vilariño-Güell et al., 2014), is involved in endocytosis of AMPARs (Perrett et al., 2015). In sum, these data reinforce AMPAR endocytosis and recycling pathways as pathways that when dysfunctional, contribute directly to neurological disorders. Our findings are a stepping stone in the understanding of molecular interactors for AMPARs, provide further evidence for the role of dysregulation of AMPAR trafficking in disease, and establish a framework for future AMPAR studies.

## Materials and Methods

### Purification of ATVs

To isolate ATVs, we followed a previously developed protocol for synaptosome generation and synaptic vesicle isolation (Ahmed et al., 2013) and extensively modified it to specifically purify ATVs. 8-12 whole mouse brains were removed from ~P20 CD-1 mice and immediately homogenized. (See Figure 1 for full summary.) This initial homogenate was spun in a JA-20 rotor at 2700 RPM (880 G) for 10 minutes to pellet blood vessels and other large cellular debris. The supernatant was then spun at 10,000 RPM (12,064 G) for 15 minutes to pellet synaptosomes. The supernatant was discarded and the periphery of the pellet was resuspended, which helps to remove mitochondria, before spinning at 11,000 RPM (14,597 G) for 15 minutes. The supernatant was again discarded, and the pellet resuspended to 5 ml total volume. The suspension was added to a Dounce homogenizer along with 45 ml of ultrapure water and was briefly homogenized to hypoosmotically lyse the synaptosomes. Immediately afterwards, 60 ul of 1 mg/ml pepstatin A and 120 ul of 200 mM PMSF in 1M HEPES was added. This solution was spun at 19,500 RPM (45,871 G) for 20 minutes to pellet plasma membrane and large cellular debris while leaving small organelles like vesicles in solution. The supernatant was then removed and spun in a Ti-70 ultracentrifuge at 50,000 RPM (256,631 G) for 2 hours at 4C to pellet small organelles like trafficking vesicles. This pellet (LP2 for “lysis pellet 2”) was transferred to a small homogenizer and resuspended in 2 ml of PBS by homogenization and mechanically sheared through a 27-gauge needle. The concentration of LP2 was checked using BCA and aliquoted into 2 mg aliquots at approximately 5 μg/μl. Any LP2 not used immediately for ATV isolation was flash frozen with liquid nitrogen and stored at −80 °C until use.

To isolate ATVs from LP2, 1 aliquot of LP2 was diluted to 1 ml total volume in 0.5% BSA in PBS, 5 ul of mouse anti-GluA1 monoclonal antibody (1ug/μl, Synaptic Systems, Gottingen, Germany) was added and allowed to bind while rotating for 12 hours at 4 °C. To prevent nonspecific binding, 50 μl of paramagnetic protein G beads (Dynabeads, ThermoFisher Scientific, Waltham, MA) were washed 3 times in 0.5% BSA in PBS for 15 minutes on ice and then 3 times in PBS for 5-minute washes on ice prior to addition of LP2. The LP2 mixture was then added to the beads and rotated for 2 hours at 4 °C. Dynabeads were separated from solution using a magnet, and the flow through was collected for Western blot analysis. ATVs were then gently eluted with three, 20-minute washes with 33 ul of GluA1 peptide (20 ug/ul) representing the antibody epitope (GenScript Biotech, Piscataway, NJ). ATVs were then immediately used and continually stored on ice at 4 °C.

### GluA1 Knockout Mice

Knockout mutant mice for *GRIA1*, the gene encoding GluA1, have been previously described (Zamanillo et al., 1999). Knockout mice were generated by interbreeding heterozygous mice.

### Western blots

For Western blot analysis, samples were first separated by SDS-PAGE and then electrophoretically transferred onto membranes. After transfer, the membranes were then treated with blocking buffer and labeled using an iBind Flex (ThermoFisher Scientific). GluA1 (Abcam, Cambridge, UK), GluN1 (Synaptic Systems), PSD-95 (Abcam), VGLUT1 (Abcam), Lamp1 (Proteintech, Rosemont, IL), and golgin (Abcam) were each individually probed. The bands were visualized either by immunofluorescence with a LI-COR Odyssey (Lincoln, NE) or with chemiluminescence with a Konica Minolta - SRX101A (Tokyo, Japan).

### Transmission electron microscopy

Negative stained transmission electron microscopy (TEM) was performed on ATVs. Copper mesh grids were glow discharged in argon gas for 20 seconds before 4 ul of ATV eluate was applied and allowed to settle for 30 minutes. The grid was then washed three times with ultra-pure water. The grid was negatively stained using 1% uranyl acetate for 2 min then blotted and allowed to dry at room temperature for 20 minutes. The grid was imaged using a JEOL 1400 TEM at 120 kEV. The diameters of ATVs were measured using ImageJ. Immunogold labeling was performed for GluA2 (BioLegend, San Diego, CA), GluA3 (Synaptic Systems), Syb2 (Abcam), Syt1 (Abcam), TfR (ThermoFisher Scientific), and Syp1 (Synaptic Systems). For immunogold labeling, the same protocol for negative stained TEM was performed; however, after ATV addition, the grids were incubated in a 1:50 dilution of rabbit polyclonal primary antibody in blocking buffer (0.5% BSA, 0.5% ovalbumin in PBS) for 1 hour. Then three, 5-minute washes in PBST were performed followed by a 1-hour incubation in 1:50 10 nm gold anti-rabbit secondary antibody (ThermoFisher Scientific). Three more 5-minute washes in PBST were performed, and then samples were fixed in 8% glutaraldehyde for 30 seconds. Staining and imaging were performed as previously described.

### Liquid chromatography mass spectrometry

Purified ATVs were resuspended in 50 ul 0.2 % Rapigest (Waters, Milford, MA) in 20 mM NH4HCO3 in 0.65 ml low protein binding polypropylene tubes before the addition of 5 mM DTT and incubation at 60°C for 30 min. After this, iodoacetamide was added to a final concentration of 7.5 mM and samples were incubated for 30 additional minutes. Samples were then digested with 2.5 micrograms of sequencing grade trypsin (Trypsin Gold, Mass spectrometry grade, Promega, Madison, WI) at 37 °C, overnight. A second aliquot of trypsin (1.5 ug) was added, and the samples incubated for an additional 3 hours at 37 °C. After this, samples were acidified by adding 5% formic acid and incubated for 30 minutes at room temperature. Tryptic peptides were recovered from the supernatant by C18 solid phase extraction using ZipTips (MilliporeSigma, Burlington, MA), eluted in two, 7 ul drops of 50% acetonitrile and 0.1% formic acid, and evaporated and resuspended in 5 ul 0.1% formic acid for LC-MS/MS analysis.

Peptides resulting from trypsinization were analyzed on a QExactive Plus mass spectrometer (ThermoFisher Scientific) connected to a NanoAcquity™ Ultra Performance UPLC system (Waters). A 15-cm EasySpray C18 column (ThermoFisher Scientific) was used to resolve peptides (60-min 2–30% B gradient with 0.1% formic acid in water as mobile phase A and 0.1% formic acid in acetonitrile as mobile phase B, at a flow rate of 300 nl/min). MS was operated in data-dependent mode to automatically switch between MS and MS/MS. MS spectra were acquired between 350 and 1500 m/z with a resolution of 70000. For each MS spectrum, the top ten precursor ions with a charge state of 2+ or higher were fragmented by higher-energy collision dissociation. A dynamic exclusion window was applied which prevented the same m/z from being selected for 10 seconds after its acquisition.

Peak lists were generated using PAVA in-house software (Guan et al., 2011). All generated peak lists were searched against the mouse subset of the UniProtKB database (SwissProt.2013.6.17) (plus the corresponding randomized sequences to calculate false discovery rate on the searches), using Protein Prospector (Clauser et al., 1999). The database search was performed with the following parameters: a mass tolerance of 20 ppm for precursor masses and 30 ppm for MS/MS, cysteine carbamidomethylation as a fixed modification, and acetylation of the N terminus of the protein, pyroglutamate formation from N terminal glutamine, and oxidation of methionine as variable modifications. A 1% false discovery rate was permitted at the protein and peptide level. All spectra identified as matches to peptides of a given protein were reported, and the number of spectra (peptide spectral matches, PSMs) was used for label free quantitation of protein abundance in the samples. Abundance index for each protein was calculated as the ratio of PSMs for a protein to the total PSMs for all components identified in the run divided by the polypeptide molecular weight.

## Acknowledgements

We thank Lu Chen, Robert Malenka, and Richard Held for discussions, Robert Malenka for kindly providing the GluA1 knockout mice, and Thomas Südhof and Lu Chen for kindly providing wildtype mice. Additionally, we thank the National Institutes of Health (grant R37MH063105 to ATB) and the National Science Foundation Graduate Research Fellowship (grant 2016205587 to JJP) for support. Mass spectrometry was provided by the Mass Spectrometry Resource at UCSF (A.L. Burlingame, Director) supported by the Dr. Miriam and Sheldon G. Adelson Medical Research Foundation (AMRF) and NIH P41GM103481 and 1S10OD016229. The project described was supported, in part, by ARRA Award Number 1S10RR026780-01 from the National Center for Research Resources (NCRR). Its contents are solely the responsibility of the authors and do not necessarily represent the official views of the NCRR or the National Institutes of Health.

---

**Supplemental Figure 1.**
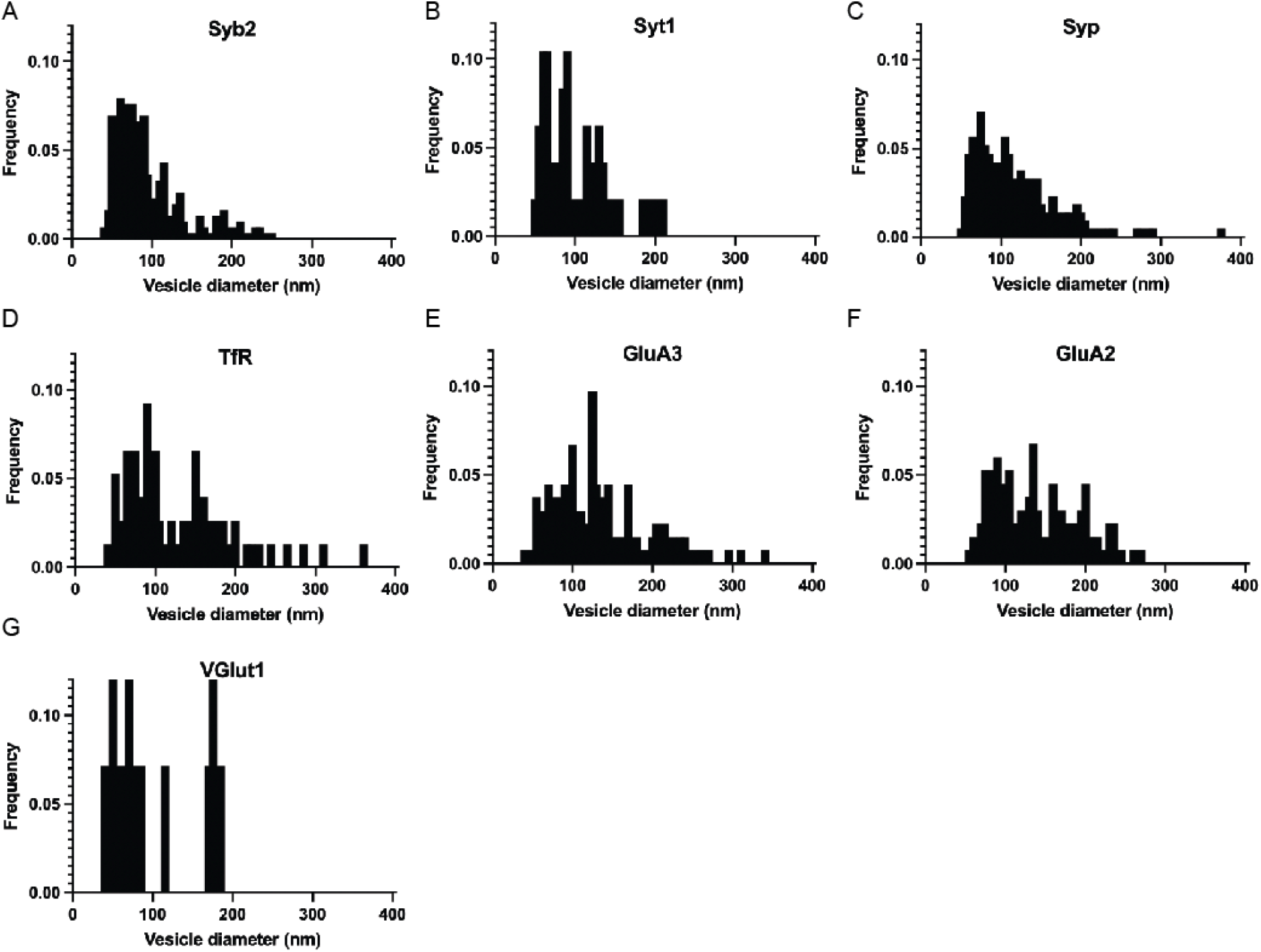
Normalized frequency distribution of diameters of ATVs labelled during immuno EM Normalized frequency distribution of diameters of ATVs labelled with **(A)** Syb2 **(B)** Syt1 **(C)** Syp **(D)** TfR **(E)** GluA3 **(F)** GluA2 **(G)** VGlut1.

## Source Data Figure Legends

*Source Data Figure 1B.*

**(A)** Unaltered Western blots. **(B)** Western blots with lanes labelled and relevant bands labelled (green arrows).

*Source Data Figure 1F.*

**(A & C)** Unaltered Western blots. **(B & D)** Western blots with lanes labelled and relevant bands labelled (green arrows).

